## Supplementary Information for "A power law of cortical adaptation"

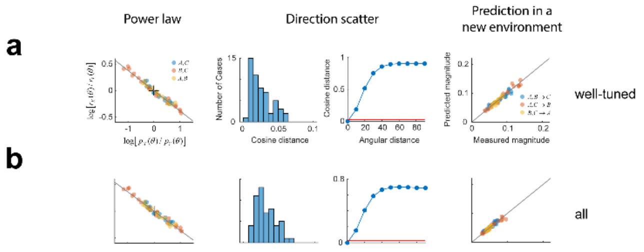


**Supplementary Figure 1.** Data selection has little impact on the results. **a,** Analysis of a dataset by selecting well-tuned neurons with circular variance less than 0.5, and **b,** Analysis of the same dataset using all neurons. Only minor differences are observed. The asymptotic cosine distance between response vectors between two angles is lower when we include all neurons. The equivalent angular difference increases as well. Both the power law and its prediction remain largely unchanged.

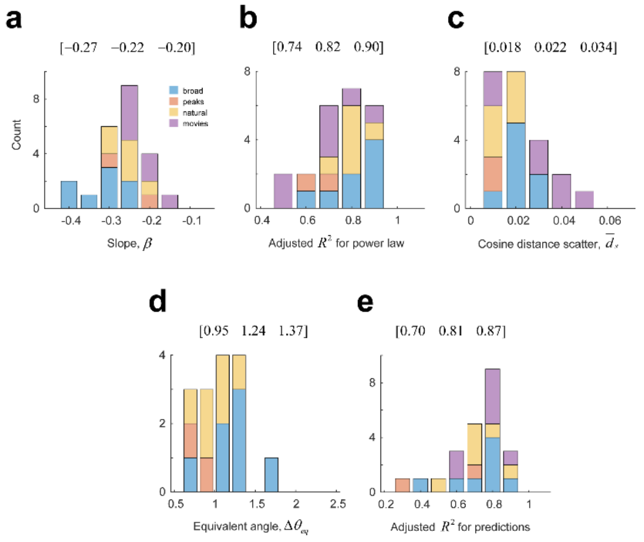


**Supplementary Figure 2**. Distribution of data fits**. a,** Distribution of the slope in the power law across all our experiments. Different experiment types are assigned different colors. Slopes were largest for experiments using broad von Mises distributions and lowest using natural movie sequences. The numbers at the top indicate the 25th, 50th and 75th percentiles of the distribution. **b,** Distribution of the adjusted R-squared goodness of fit for the power law. **c,** Distribution of cosine distance scatter, **d,** Distribution of the equivalent angle, **e,** Distribution of R-squared values for the predictions in new environments. Note these values have almost the same distribution as in panel **b**.

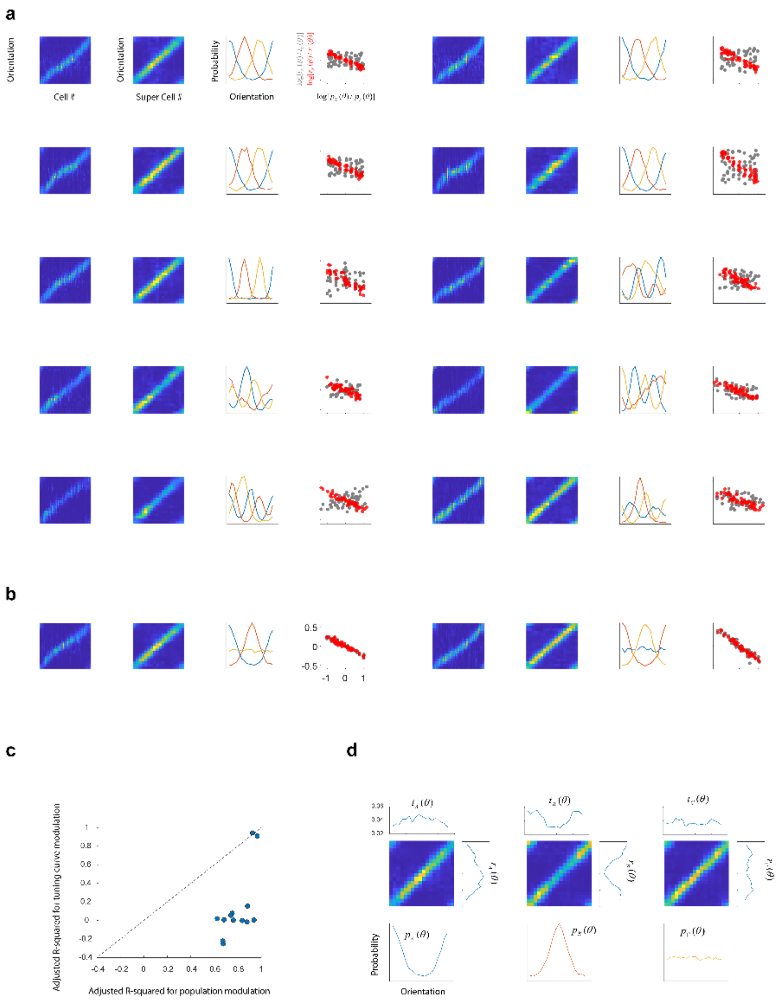


**Supplementary Figure 3**. Assessing the contributions of modulations in population versus modulations in tuning curves. These are analogous to the gain factors in a prior study[31]. **a**, Each set of four panels illustrates the analysis for a separate experiment. First, for each environment, we convert the matrix representing the responses for each cell, shown in the leftmost panel, into one where neurons are grouped by their preferred orientation in 18 different bins and the results averaged. Thus, the second panel in the row is a matrix of size 18 x 18 (18 orientations and 18 "super neurons"). The $l_2$ norms of the rows in this matrix are the population norms $r_X(\theta)$, where $\theta$ is the orientation of a stimulus. Similarly, we can take the $l_2$ norms of the columns, which represents the tuning curve norms, which we denote by $t_X(\theta)$. Here, $\theta$ represents the preferred orientation of a super-neuron. Suppose we adapt the population to an orientation $\theta$. Is the change in the population best described by a change in $t_X(\theta)$ or a change in $r_X(\theta)$? More generally, do the ratios of the population magnitudes or the ratios of tuning curve magnitudes best correlate with the ratios of stimulus priors? The scatter plots show that the population magnitudes do a much better job than the tuning curve magnitudes. **b,** The only instances where the tuning curve magnitudes provided as good a prediction those of the population were in situations where the environments were simple von Mises distributions 90 deg apart with a uniform distribution. **c,** Across our experiments, the adjusted R-squared values are substantially higher when the population magnitudes are used compared to those of tuning curve magnitudes (except for the two isolated cases mentioned in **b**). **d**, The reasons that both population and tuning curve magnitudes are approximately the same in the simple environments shown in **b**, are the following. First, the matrix is nearly diagonal for super-cells that are well tuned. Second, the environments cause a modulation of responses along the diagonal. Third, this means that the norms of columns and rows are approximately the same. The effect along the diagonal, therefore, is not well-defined. Different combinations of population and tuning curve modulations can explain it. These analyses suggest that complex environments are better suited at discriminating between modulation of the population and modulation of the tuning curves. In our data, the former explanation prevails.

| animal | type | sex | ncells | selected | gOSI 25 | gOSI 50 | gOSI 75 | equiv_angle | mean_dth | beta | R2_pl | R2_pr | R2_pl_o | R2_pr_o | kopt | norm_a | R2_norm |
|---|---|---|---|---|---|---|---|---|---|---|---|---|---|---|---|---|---|
| adpt08_001_000 | 1 | F | 913 | 443 | 0.282 | 0.494 | 0.737 | 0.897 | 0.016 | -0.251 | 0.930 | 0.801 | 0.944 | 0.813 | 3.360 | 0.507 | 0.898 |
| adpt08_002_000 | 1 | F | 906 | 263 | 0.196 | 0.342 | 0.567 | 1.185 | 0.021 | -0.291 | 0.897 | 0.803 | 0.899 | 0.808 | 23.357 | 0.522 | 0.925 |
| adpt11_001_000 | 1 | M | 767 | 205 | 0.189 | 0.328 | 0.543 | 1.379 | 0.026 | -0.380 | 0.972 | 0.945 | 0.978 | 0.963 | 8.859 | 0.504 | 0.936 |
| adpt08_011_000 | 1 | F | 1001 | 304 | 0.231 | 0.377 | 0.582 | 1.220 | 0.020 | -0.220 | 0.948 | 0.889 | 0.949 | 0.895 | 18.330 | 0.543 | 0.909 |
| adpt08_012_000 | 1 | F | 689 | 161 | 0.165 | 0.283 | 0.489 | 1.877 | 0.030 | -0.259 | 0.885 | 0.754 | 0.890 | 0.773 | 11.288 | 0.545 | 0.895 |
| adpt09_070_000 | 1 | F | 856 | 275 | 0.206 | 0.351 | 0.607 | 1.480 | 0.024 | -0.240 | 0.772 | 0.601 | 0.775 | 0.608 | 18.330 | 0.528 | 0.928 |
| adpt09_071_000 | 1 | F | 675 | 124 | 0.161 | 0.277 | 0.457 | 2.369 | 0.035 | -0.365 | 0.693 | 0.488 | 0.693 | 0.488 | 100.000 | 0.544 | 0.935 |
| adpt09_081_000 | 1 | F | 724 | 186 | 0.181 | 0.313 | 0.528 | 1.342 | 0.023 | -0.307 | 0.865 | 0.689 | 0.902 | 0.817 | 4.281 | 0.516 | 0.933 |
| adpt09_100_000 | 2 | F | 764 | 288 | 0.211 | 0.385 | 0.639 | 0.774 | 0.014 | -0.169 | 0.249 | -0.211 | 0.789 | 0.729 | 2.637 | 0.497 | 0.840 |
| adpt09_101_000 | 2 | F | 650 | 150 | 0.160 | 0.292 | 0.493 | 1.020 | 0.018 | -0.251 | 0.219 | -0.694 | 0.676 | 0.381 | 4.281 | 0.484 | 0.868 |
| adpt10_010_000 | 3 | F | 534 | 149 | 0.190 | 0.309 | 0.560 | 0.955 | 0.017 | -0.255 | 0.785 | 0.547 | 0.831 | 0.713 | 4.281 | 0.498 | 0.764 |
| adpt10_050_000 | 3 | F | 500 | 151 | 0.168 | 0.310 | 0.590 | 0.800 | 0.014 | -0.203 | 0.716 | 0.639 | 0.818 | 0.772 | 2.637 | 0.512 | 0.944 |
| adpt10_060_000 | 3 | F | 501 | 134 | 0.164 | 0.294 | 0.533 | 0.949 | 0.018 | -0.200 | 0.753 | 0.526 | 0.821 | 0.568 | 3.360 | 0.486 | 0.760 |
| adpt09_130_000 | 3 | F | 644 | 177 | 0.197 | 0.337 | 0.558 | 1.357 | 0.022 | -0.258 | 0.922 | 0.920 | 0.928 | 0.932 | 18.330 | 0.463 | 0.844 |
| adpt09_150_000 | 3 | F | 984 | 239 | 0.153 | 0.272 | 0.495 | 1.257 | 0.022 | -0.224 | 0.701 | 0.503 | 0.830 | 0.707 | 3.360 | 0.487 | 0.883 |
| adpt11_011_000 | 3 | M | 459 | 118 | 0.188 | 0.317 | 0.522 | 1.264 | 0.022 | -0.187 | 0.738 | 0.820 | 0.754 | 0.722 | 1.000 | 0.522 | 0.962 |
| adpt09_500_000 | 4 | F | 1287 | 1287 | | | | | 0.043 | -0.204 | 0.808 | 0.876 | | | | 0.615 | 0.931 |
| adpt09_501_000 | 4 | F | 1286 | 1286 | | | | | 0.053 | -0.198 | 0.582 | 0.634 | | | | 0.676 | 0.875 |
| adpt10_200_000 | 4 | F | 657 | 657 | | | | | 0.039 | -0.206 | 0.705 | 0.857 | | | | 0.679 | 0.935 |
| adpt10_500_000 | 4 | F | 1379 | 1379 | | | | | 0.036 | -0.210 | 0.731 | 0.699 | | | | 0.622 | 0.900 |
| adpt10_700_000 | 4 | F | 457 | 457 | | | | | 0.017 | -0.218 | 0.903 | 0.900 | | | | 0.429 | 0.829 |
| adpt11_012_000 | 4 | M | 490 | 490 | | | | | 0.050 | -0.133 | 0.599 | 0.826 | | | | 0.701 | 0.647 |
| adpt11_030_000 | 4 | M | 601 | 601 | | | | | 0.016 | -0.178 | 0.770 | 0.937 | | | | 0.496 | 0.871 |

**Supplementary Table 1.** Summary of experiments. Each row summarizes a different experiment. From left to right, the columns include the following information: experiment name, sex, type of experiment (codes: 1, von Mises and uniform distributions; 2, peaked distributions; 3, natural orientation distributions; 4, movie sequences), total number of cells recorded, total number of cells selected (with $\mathrm{gOSI} > 0.5$), 25$^{th}$ percentile of gOSI, 50$^{th}$ percentile of gOSI, 75$^{th}$ percentile of gOSI, equivalent angle, mean cosine distance scatter, $\beta$ fit to the power law, adjusted R-squared value for the power law relationship, adjusted R-squared value for the prediction, adjusted R-squared value for the power law relationship after smoothing correction, adjusted R-squared value for the prediction after smoothing correction, optimal $\kappa$, constant of proportionality between $l_1$ and $l_2$ norms, adjusted R-squared value for the linear fit between the norms.
